# Knockout of *Babesia bovis rad51* ortholog and its complementation by expression from the BbACc3 artificial chromosome platform

**DOI:** 10.1101/606590

**Authors:** Erin A. Mack, Yu-Ping Xiao, David R. Allred

## Abstract

*Babesia bovis* establishes persistent infections of long duration in cattle, despite the development of effective anti-disease immunity. One mechanism used by the parasite to achieve persistence is rapid antigenic variation of the VESA1 cytoadhesion ligand through segmental gene conversion (SGC), a phenomenon thought to be a form of homologous recombination (HR). To begin investigation of the enzymatic basis for SGC we initially identified and knocked out the Bb*rad51* gene encoding the *B. bovis* Rad51 ortholog. BbRad51 was found to be non-essential for in vitro growth of asexual-stage parasites. However, its loss resulted in hypersensitivity to methylmethane sulfonate (MMS) and an apparent defect in HR. This defect rendered attempts to complement the knockout phenotype by reinsertion of the Bb*rad51* gene into the genome unsuccessful. To circumvent this difficulty, we constructed an artificial chromosome, BbACc3, into which the complete Bb*rad51* locus was inserted, for expression of BbRad51 under regulation by autologous elements. Maintenance of BbACc3 makes use of centromeric sequences from chromosome 3 and telomeric ends from chromosome 1 of the *B. bovis* C9.1 line. A selection cassette employing human dihydrofolate reductase enables recovery of transformants by selection with pyrimethamine. We demonstrate that the BbACc3 platform is stably maintained once established, assembles nucleosomes to form native chromatin, and expands in telomere length over time. Significantly, the MMS-sensitivity phenotype observed in the absence of Bb*rad51* was successfully complemented at essentially normal levels. We provide cautionary evidence, however, that in HR-competent parasites BbACc3 can recombine with native chromosomes, potentially resulting in crossover. We propose that, under certain circumstances this platform can provide a useful alternative for the genetic manipulation of this group of parasites, particularly when regulated gene expression under the control of autologous elements may be important.

## Introduction

Babesiosis is a tick-borne disease caused by apicomplexan parasites of the genus *Babesia*. Humans are not the natural host for any babesial parasite but may be an incidental host, acquiring zoonotic infections with a variety of different species. *Babesia microti* is the most common species of *Babesia* to infect humans, although in western Europe infections commonly occur with *Babesia divergens*. In the U.S. infections have been observed with *Babesia duncani* and *B. divergens*-like organisms, as well as the unspeciated WA1 and MO1 isolates (reviewed in [1]). Many individuals may carry asymptomatic infections [2], including as a result of inadequate drug treatment of acute parasitemia [3, 4], posing a serious risk to the blood supply [5]. In cattle babesiosis may be caused by at least five different species, with *Babesia bovis* generally considered the most virulent. *B. bovis* shares many parallels with the human malarial parasite, *Plasmodium falciparum*, including immune evasion via cytoadhesion and antigenic variation, and the capacity for development of a lethal cerebral disease [6].

Bovine babesiosis caused by *B. bovis* is quite severe. Cattle that survive the acute disease develop a strong anti-disease immunity, but remain persistently infected for periods of at least several years. This parasite makes use of at least two mechanisms to effect persistence: (i) cytoadhesion of infected red blood cells (iRBCs) to capillary and post-capillary venous endothelium, presumably in order to avoid splenic clearance; and (ii) rapid antigenic variation of the cytoadhesion ligand, VESA1, to avoid antibody-mediated forced re-entry into the circulation [7]. Cytoadhesion is therefore a behavior that is immunologically sensitive, and which the parasite correspondingly has evolved means to protect. The development of the ability to abrogate antigenic variation would allow for ready elimination of the parasite; this capability is a desirable goal for parasite control through disruption of an essential aspect of its biology.

There is a great need to understand the molecular bases of virulence mechanisms in parasites. One informative way to facilitate such studies is through the ability to genetically manipulate the organisms. To date, the tools developed for the genetic manipulation of *B. bovis* are limited and remain most appropriate for use in characterizing individual targets, although there is a need for high-throughput methodologies [8]. The expression of exogenous genes has been demonstrated, using babesial promoter sequences to drive transcription of the target sequences. This has been achieved transiently through transfection with circular plasmids [9–13]. The longevity of this class of vector has been improved by the inclusion of a centromeric segment, improving the segregation of the circular molecules [12]. Long-term, stable expression has been achieved by single-crossover insertional mutagenesis of chromosomal loci [12, 14]. This approach demonstrates the capacity for creating targeted gene disruptions and the potential for the use of promoter trapping strategies. Significantly, double-crossover knockout of the thioredoxin peroxidase (*tpx*) gene has been achieved, providing complete gene knockouts [12]. Complementation of *tpx* was achieved by re-integration of the gene ectopically into the EF1α locus, resulting in high level expression. However, ectopic expression of exogenous genes in this manner may suffer from improper expression due to local chromatin modifications, including silencing or significant overexpression, compromising interpretation of genetic contributions to phenotype. Despite the available tools there remains a need for convenient platforms that are stable, efficiently and accurately segregated during mitosis, allow the transcription of large sequences, and which may be properly regulated relative to metabolic need.

Antigenic variation in *B. bovis* is known to heavily utilize segmental gene conversion (SGC) to modify the VESA1 ligand (BAK), and may participate in in situ transcriptional switching (isTS) as well. SGC is thought to be a form of homologous recombination (HR), much the same as gene conversion, a process that is dependent upon Rad51 proteins. To initiate investigation of the enzymatic underpinnings of antigenic variation by SGC in *B. bovis*, we describe here the knockout of the *B. bovis rad51* gene by double crossover replacement [15]. Given the potential for adverse effects if expressed at inappropriate levels, we did not want to overexpress Bb*rad51* during complementation. Moreover, because of the knockout of the Bb*rad51* gene, homologous recombination was lost and complementation of the knockout by reintroduction of the gene into the genome was not possible. We have addressed this issue through artificial chromosome technology, an approach that allows the complementation of disrupted genes by reintroduction of the entire locus, providing gene control through its autologous regulatory elements. An artificial chromosome platform has already been developed for *Plasmodium berghei*, enabling transformation efficient enough for shotgun screening to recover genes associated with drug resistance [16, 17]. We have created an analogous platform for use with *B. bovis* through the assembly of an artificial chromosome, BbACc3. BbACc3 is comprised of *B. bovis* centromeric and telomeric ends in a modified pBluescript-KS(+) backbone. This construct employs the human dihydrofolate reductase (h*DHFR*) gene as selectable marker, conferring resistance to pyrimethamine. BbACc3 was constructed and is manipulated as the circular plasmid, pBACc3, which is linearized for transfection. Linearization with PmeI removes a spacer sequence and exposes legitimate telomeric ends. Here, we demonstrate that BbACc3 transforms *B. bovis* to pyrimethamine resistance, is maintained as a distinct linear chromosome that assembles into chromatin, and undergoes expansion of its telomeres. Significantly, the Bb*rad51* gene expressed from BbACc3 under regulation by its autologous elements provided normal levels of complementation of the Bb*rad51* knockout upon challenge with methylmethane sulfonate (MMS). Although further optimization is desirable, this platform may facilitate long-term complementation of essential genes for studies connecting genotype and phenotype, particularly where improper regulation would be problematic.

## Results

### The orthologous Bbrad51 gene was identified

The *B. bovis rad51* gene (Bb*rad51*) was identified by first searching the genus *Babesia* non-redundant proteome (S1 Table). The search was initiated with the *Saccharomyces cerevisiae* Rad51 sequence (CAA45563) as bait, using BLAST, with default parameters [18]. In this initial search, proteins XP_001609877 (E= 2e-103) and XP_001609660 (E= 4e-15) were identified as candidates. To cast a wider net and ensure capture of all candidates, CAA45563 was again used to initiate an iterative search of the genus *Babesia* non-redundant protein database, using Psi-BLAST [19, 20] with default parameters [19]. After three iterations the top *B. bovis* candidate, XP_001609877, had a significance score of E = 3e-156, and three additional proteins of the Rad51/DMC1/RadA superfamily were also identified. Each of these proteins then was used individually in reciprocal iterative searches of the NCBI non-redundant protein database, excluding the genus *Babesia*, using Delta-BLAST and composition-based statistics to provide their presumptive identities [19, 20]. These proteins included the second-best fit, XP_001609660 (E = 3e-86; XRCC3-like), XP_001610815 (E= 8e-16; Rad51 homolog 2-like), and XP_001609995 (E= 0.077; XRCC2-like). Genes encoding putative orthologs of all four proteins are found among various other piroplasms (S1 Table; adapted from PiroplasmaDB). For XP_001609877 this search yielded over 200 significant hits of E= 0.0, all of which were confirmed or putative eukaryotic Rad51 molecules sharing 47-49% identity. Alignment of the four candidates with a number of confirmed or predicted Rad51 proteins demonstrated excellent alignment of all functional domains and motifs (S1 Fig.). An alignment tree was created by the Neighbor-joining method, using Jukes-Cantor protein distance measures and 100 bootstraps to assess robustness. The tree shows that, among the apicomplexan parasites Rad51 proteins cluster in apparent clades (Fig. 1A, colored blocks), suggesting shared selection pressures within individual groups. *P. knowlesi* was the only exception, failing to cluster with other *Plasmodium spp*. XP_001609877 clusters with other eukaryotic Rad51 proteins, forming an apparent clade with other piroplasmids, yet is clearly quite different. By contrast, the other three *B. bovis* RecA/RadA/Rad51-related proteins cluster more closely with the *Escherichia coli* RecA protein included as outlier. An analysis of XP_001609877 for probable structural domains by the Conserved Domains Database yielded specific hits of PTZ00035 (“Rad51 protein, provisional”; residues 6-343; E= 0e+00) and cd01123 (“Rad51_DMC1_radA, P-loop NTPase superfamily”; residues 105-339; E= 3.33e-111), as well as a lower-confidence hit in the N-terminus for pfam14520, a small “helix-hairpin-helix” domain (residues 39-83; E= 2.02e-03). The pfam14520 “helix-hairpin-helix” domain, which is common to Rad51 proteins, was lacking from the other three candidates. Further support for the putative assignment of this protein as a legitimate Rad51 ortholog was obtained by ab initio prediction of its three-dimensional structure, using Robetta software (http://robetta.bakerlab.org/) [21, 22]. The predicted structure models were superimposed onto the crystal structure of the *S. cerevisiae* Rad51 H352Y mutant protein (PDB accession 3LDA) [23], using the Chimera Matchmaker algorithm [24]. The predicted three-dimensional structure of XP_001609877 (model 4) was found to coordinate spatially extremely well with the 3LDA crystal structure, including Walker A and B motifs, residue Q298 regulating access of ATP to the active site [23], and the interaction domain sequences pivotal to filament formation (Fig. 1B). Based on these results we refer to XP_001609877 as BbRad51, and BBOV_II003540 as the Bb*rad51* gene encoding this protein. The Bb*rad51* gene was annotated with two predicted introns [25], a structural model that was confirmed by RT-PCR amplification, cloning, and sequencing of the full-length transcript. Flanking sequences were recovered through 5’- and 3’-RACE reactions (see Methods). The results exactly matched the published *B. bovis* T2Bo isolate genome sequence, including predicted start and stop codons, and intron boundaries (Fig. 2A; [15, 25]). Active transcription of Bb*rad51* by asexual stages has been reported previously [26], and was confirmed here by RT-PCR (S2 Fig., panel A).

**Fig. 1.**
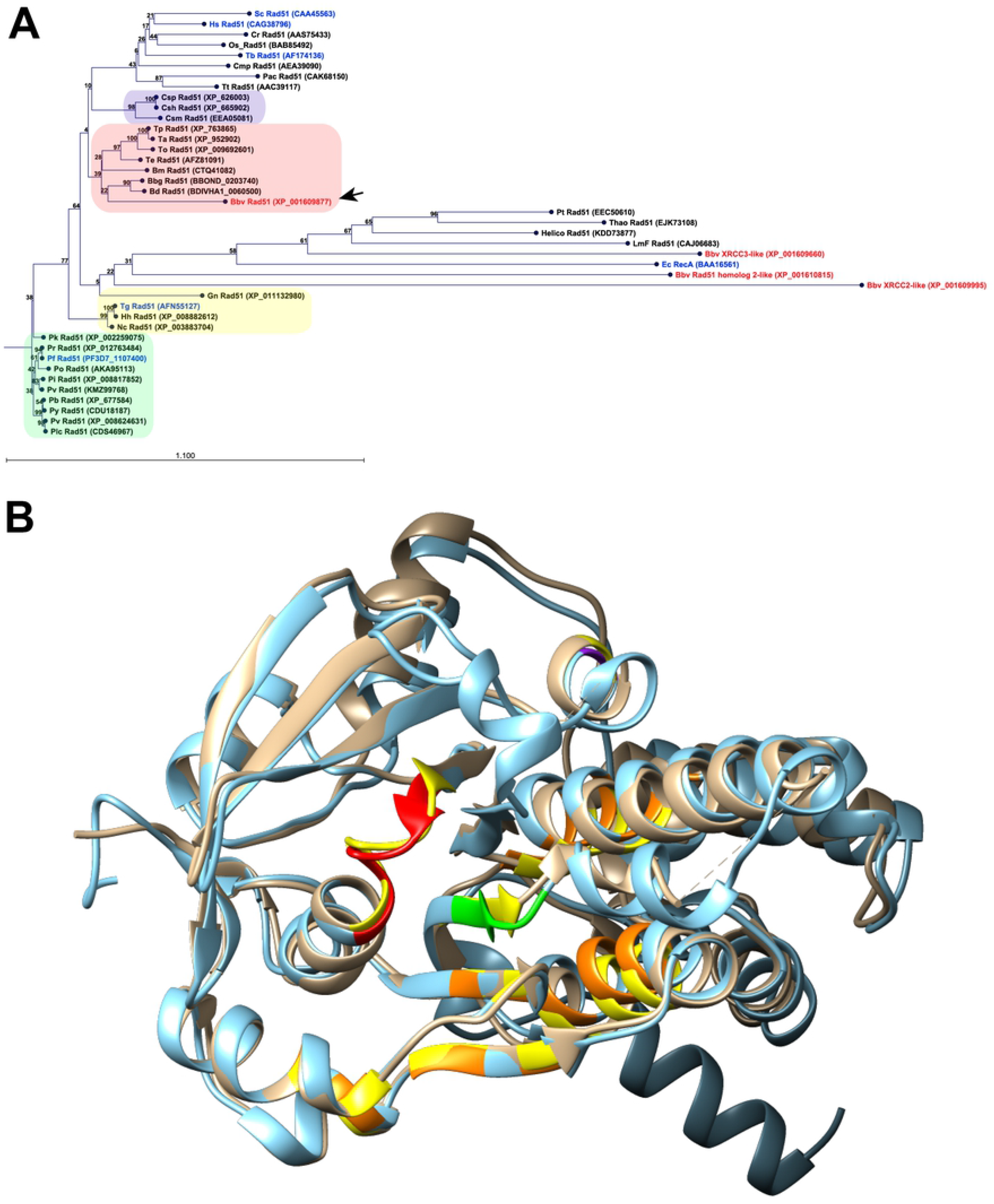
Relationship of BbRad51 to other Rad51 proteins. **A.** Alignment tree of the four *B. bovis* RecA/RadA/Rad51-related proteins (accession numbers in red) with established or predicted Rad51 proteins of other species (Neighbor-joining; Kimura protein distance measure; 100 bootstrap replicates; numbers at nodes are bootstrap values). Direct experimental evidence supports the catalysis of canonical RecA/RadA/Rad51 activities by those proteins whose accession number is indicated in blue. BbRad51 (black arrow) clusters with, but is comparatively dissimilar to, putative Rad51 proteins of other Piroplasmida (pink panel). In contrast, the three remaining *B. bovis* proteins cluster more closely with the prokaryotic (*E. coli*) RecA protein included as outlier to the alignment, and with several algal Rad51 proteins. All four *B. bovis* proteins cluster away from other apicomplexan parasites (Haemosporida, green panel; Coccidia, yellow panel) or higher eukaryotes. Interestingly, *Cryptosporidium spp*. cluster separately from other Coccidia (blue panel). **B.** The predicted three-dimensional structure of BbRad51 (blue ribbon), superimposed upon the crystal structure of the *S. cerevisiae* Rad51 H352Y mutant (3LDA [23], tan ribbon), shows excellent conformational coherence. Indicated on BbRad51 are the Walker A (red) and Walker B (green) motifs, subunit interaction domains (orange), and the position of residue Q298 that acts as “gatekeeper” controlling access of ATP to the active site (purple). Corresponding ScRad51 residues are shown in yellow. ScRad51 has a disordered N-terminal extension that fails to yield usable crystal data [71], resulting in a lack of sequence to align with that region of BbRad51.

**Fig. 2.**
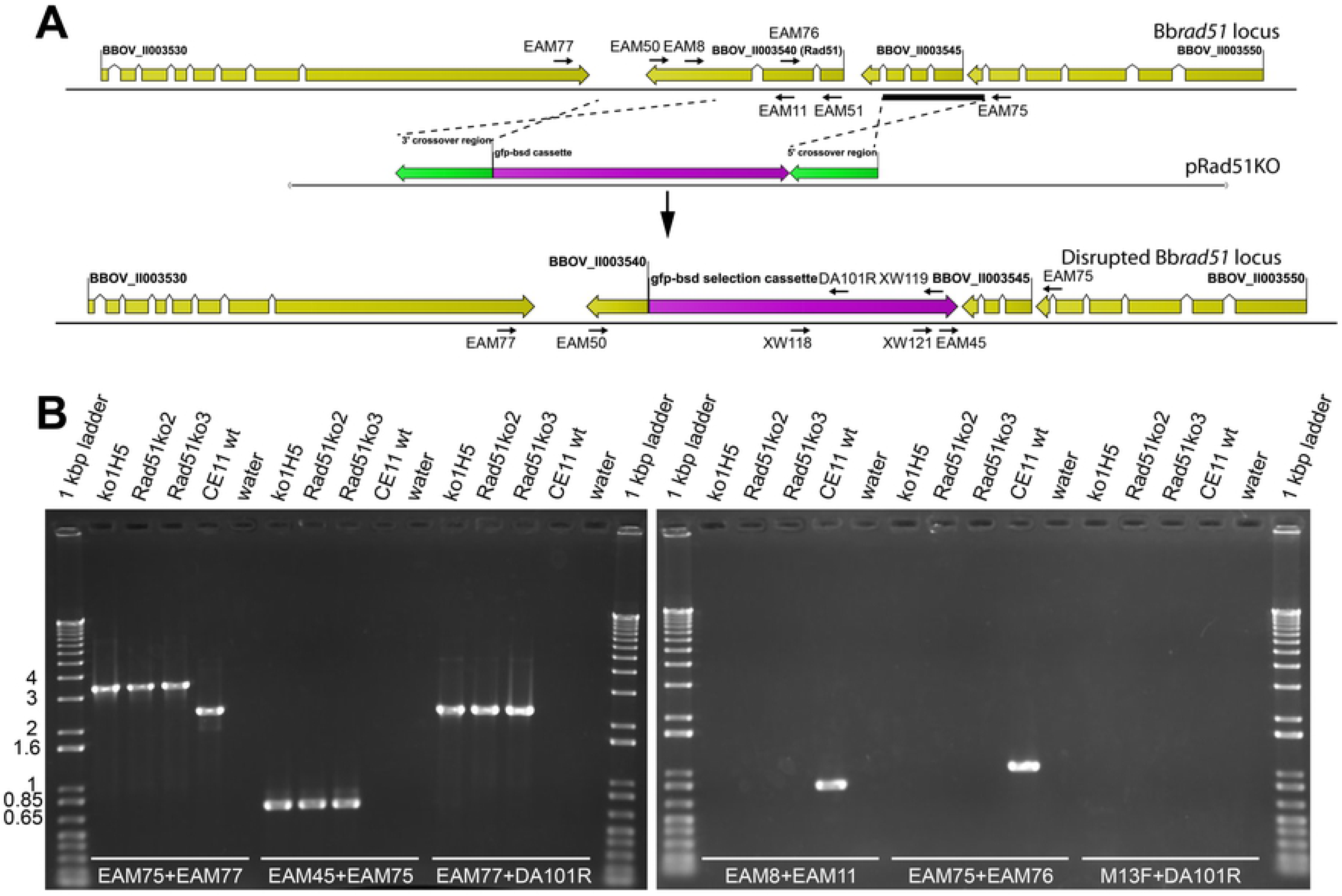
Bb*rad51* locus and knock-out strategy. **A.** The wt Bb*rad51* locus structure was confirmed to be comprised of three exons/ two introns. The gene was replaced by double homologous crossover, with a selectable *gfp-bsd* cassette (purple arrow) driven by the *B. bovis EF1*α-B hemi-promoter [10, 14], using flanking sequences from the locus (crossover regions shown as green arrows). The structures of the Bb*rad51* wt locus (top), pRAD51KO plasmid (center), and the disrupted Bb*rad51* locus (bottom) are shown. Primers used in PCR reactions and Southern blotting (S2 Table), and the sites to which each anneals, are indicated. **B.** Diagnostic PCR was used to initially confirm Bb*rad51* knock-out in each line, using the primer pairs indicated. Locus structure was fully confirmed for the ^ko1^H5 mutant line (lanes ko1H5), using PCR, Southern blotting and sequencing. Bb*rad51* knock-out was supported by diagnostic PCR for CE11Δ*rad51*^ko2^ and CE11Δ*rad51*^ko3^ (lanes Rad51ko2 and Rad51ko3, respectively).

### Bb*rad51*-null *B. bovis* lines were established

Bb*rad51* is a single copy gene in the haploid *B. bovis* genome [25]. Therefore, a single-locus double-crossover strategy was used to replace the gene with a selection cassette expressing GFP-Blasticidin-s deaminase fusion protein (*gfp-bsd*; a gift from C.E. Suarez [10, 14]). The transfection plasmid, pRad51KO (Fig. 2A), was constructed and used to transform *B. bovis* CE11 line parasites to blasticidin-s resistance. The transformed culture, and the clonal line CE11Δ*rad51*^ko1^, and subclones CE11Δ*rad51*^ko1^H5, CE11Δ*rad51*^ko1^C3, and CE11Δ*rad51*^ko1^H6 derived from it, were screened by PCR (primers used in this study are provided in S2 Table). For brevity, CE11Δ*rad51*^ko1^H5 is hereafter referred to as “^ko1^H5”. The structure of the modified Bb*rad51* locus and loss or gain of Bb*rad51* or *gfp-bsd* transcription, respectively, was confirmed by diagnostic PCR, RT-PCR, and Southern blotting, confirming the replacement of Bb*rad51* with the *gfp-bsd* selection cassette from pRad51KO (Fig. 2B; S2 and S3 Figures). PCR-mediated cloning and sequencing of the modified locus confirmed that error-free double-crossover homologous recombination (HR) had occurred. Two additional independent knockout clonal lines, CE11Δ*rad51*^ko2^ and CE11Δ*rad51*^ko3^ (for brevity, referred to as “ko2” and “ko3”, respectively), were established approximately one year later. These two lines were confirmed by diagnostic PCR and sequencing of the insertion sites (not shown). Because of the lag in obtaining the latter two lines, tests of phenotype initially were performed with ^ko1^H5 and subsequently confirmed with the latter two. As the results were generated in different experiments they are presented separately.

### Bbrad51 knockout had no effects on growth or morphology

Knockout of Bb*rad51* had no observable effects on parasite morphology at the level of light microscopy (not shown). Despite the expected significance of Rad51 function to DNA repair and genome stability, no significant differences were observed in the growth rates of wild type and Bb*rad51* knockout lines (Fig. 3). The growth data shown was obtained with a SybrGreen-based assay of cumulative DNA content to avoid the subjectivity associated with microscopic assays, although microscopic assessments of the percentage of parasitized erythrocytes on Giemsa-stained smears of cultured parasites confirmed these results (not shown). Growth was followed for 48 hours only, to avoid any artifactual differences caused by a need for media changes or cell manipulations. However, as these parasites are not developmentally synchronous in their growth, and have an 8-10 hour asexual cell cycle [27], this period represents approximately 5-6 complete cell cycles. Mean differences in growth rate of as little as 2% per generation would result in larger differences than were observed. Thus, within the limits of sensitivity of our assay no differences in growth rates were observed between wild type and knockout lines.

**Fig. 3.**
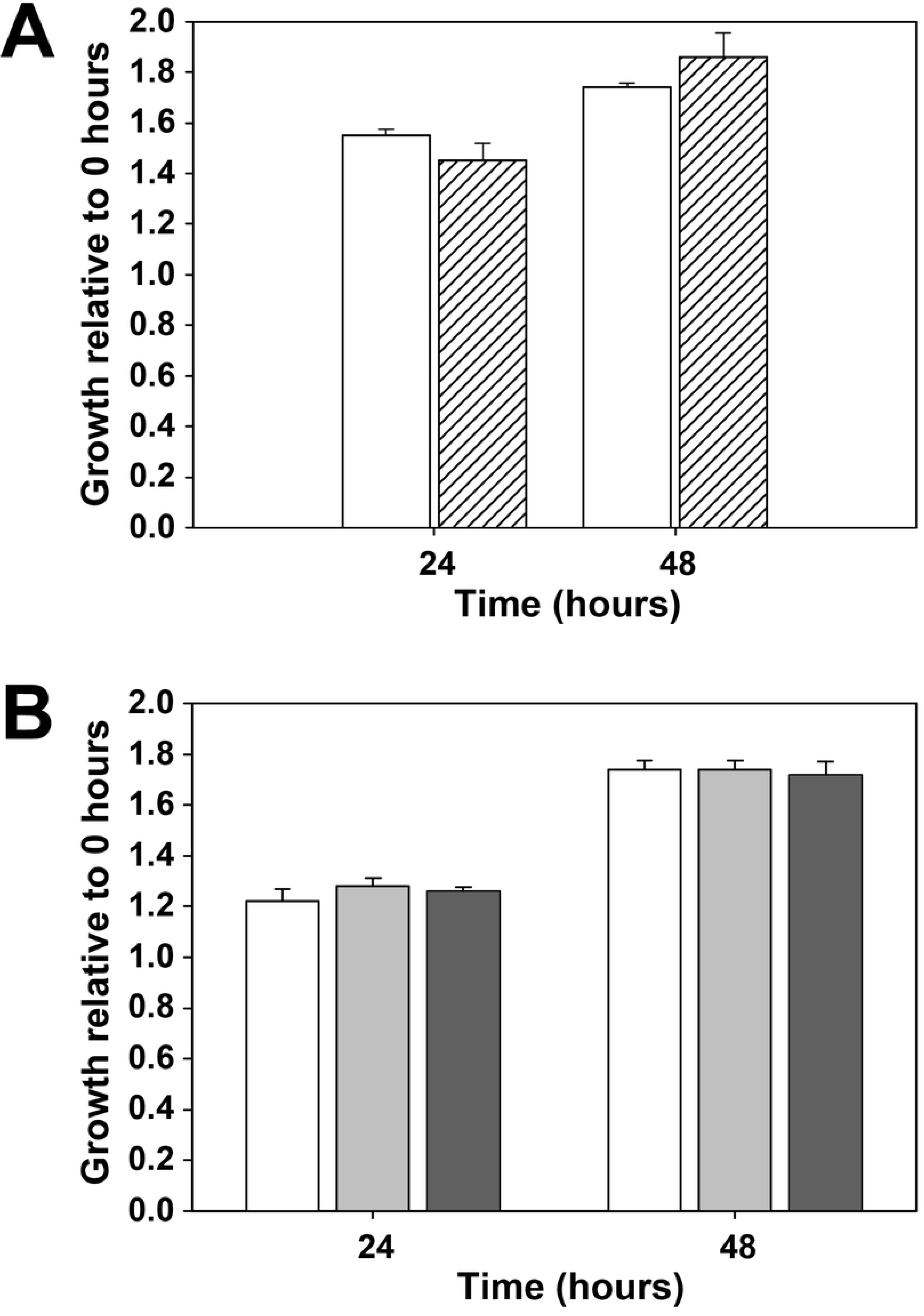
Growth of wild type and Bb*rad51* knockout parasites. Shown is the comparative growth over a 48h period (mean ± 1 s.d.) of **A.** *B. bovis* CE11 wt and the Bb*rad51* knockout clonal line, ^ko1^H5, and of **B.** CE11 wt and the CE11Δ*rad51*^ko2^ and CE11Δ*rad51*^ko3^ lines. The 48h period of the assay represents 5-6 cell cycles. The data in A. and B. were generated in independent experiments separated by a year, and so are presented separately. CE11, white bars; ^ko1^H5, hatched bars; CE11Δ*rad51*^ko2^, gray bars; CE11Δ*rad51*^ko3^, dark gray bars.

### Assessment of MMS-sensitivity

One common means of assessing contributions of a protein to DNA repair is to determine sensitivity to DNA damage deliberately induced by chemical insult. In this study, we employed exposure to methylmethane sulfonate (MMS), a chemical which alkylates guanines to 7-methylguanine and adenines to 3-methyladenine [28]. To apply that approach in this situation, sensitivity of *B. bovis* CE11 to MMS first was titrated over a range from 0 to 2000 μM, employing a strategy of exposure to MMS for 90 minutes, with a buffer washout of the MMS, followed by growth for 72 hours (S4 Fig.). Initially, we used a microscopic approach to follow parasite growth, to avoid any concerns about MMS effects on erythrocytes or residual DNA fluorescence from dead parasites. The results ranged from little or no effect at ≤250 μM to complete killing by exposure to 2000 μM MMS. *B. bovis* CE11 and ^ko1^H5 lines then were compared routinely over a range from 125- 1000 μM. *B. bovis* ^ko1^H5 was significantly more susceptible to DNA damage at 250 or 500 μM MMS than CE11 wild-type parasites (Fig. 4). In the CE11 line there was little or no effect seen at 250 μM. *B. bovis* CE11 parasites exposed to 500 μM showed a severe reduction in growth rate that quickly increased, approaching that of 0 μM controls by 48 hours. In contrast, ^ko1^H5 parasites were significantly reduced in growth at 250 μM, and 500 μM-treated parasites failed to regain normal growth even by 72 hours post-treatment. These results indicate that BbRad51 plays a role in recovery from DNA damage due to nucleotide alkylation [29–32].

**Fig. 4.**
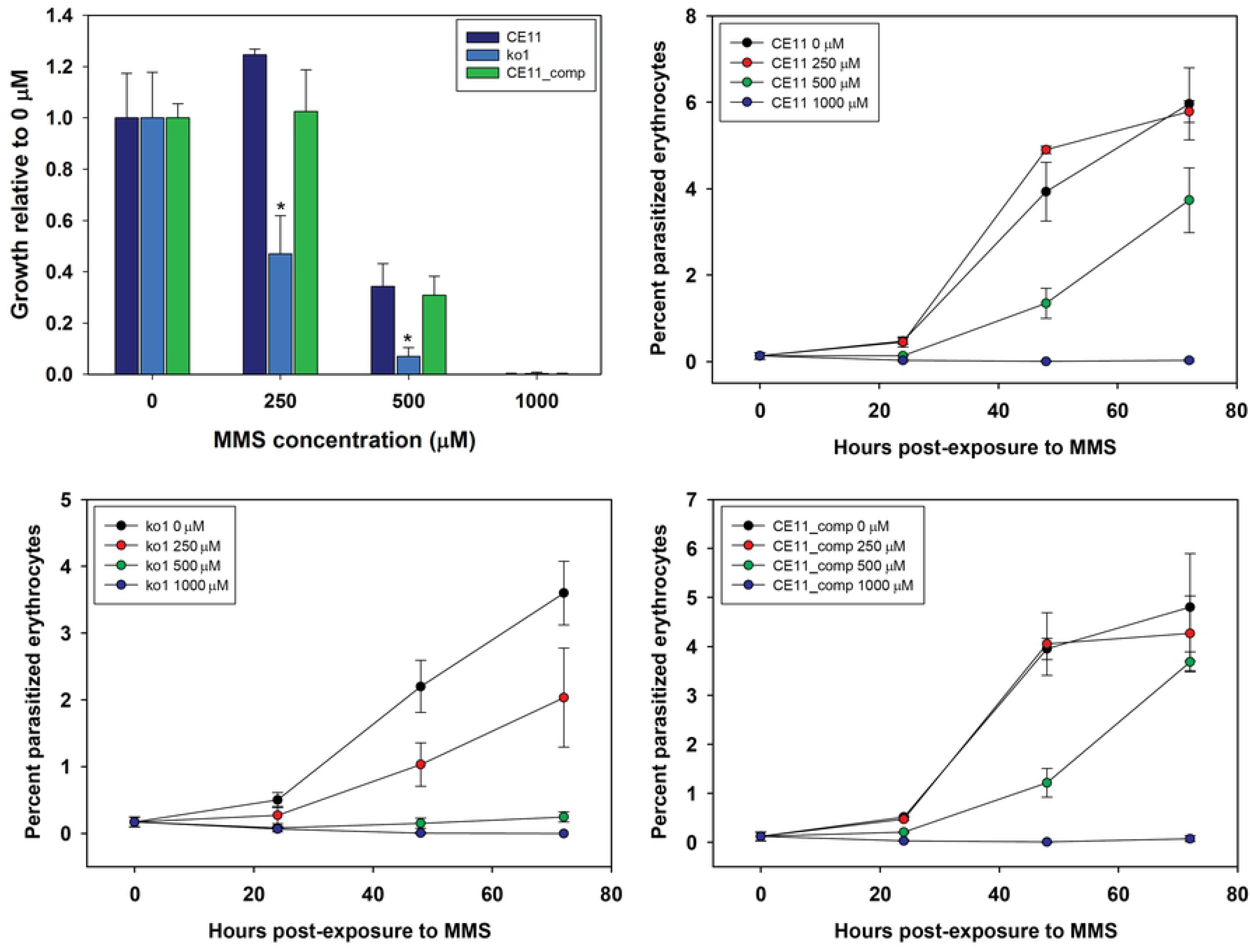
Effects of environmental insult on growth of Bb*rad51* knockouts. *B. bovis* CE11 wild-type (CE11), ^ko1^H5 (ko1), and CE11 parasites with pBb*rad51*wt_comp integrated at the Bb*rad51* locus (CE11_comp) were tested for relative sensitivity of growth to a 90-minute exposure to varying concentrations of MMS, in a 72h terminal experiment (upper left panel). Growth characteristics over time are shown for CE11 (upper right), ^ko1^H5 (lower left), and CE11_comp (lower right) following MMS exposure at the various concentrations. These results suggest that the Bb*rad51* locus is neither affected by nor refractory to manipulation by recombination.

### Re-integration of Bb*rad51* gene

To confirm the contribution of BbRad51 to this phenotype, we attempted to complement *B. bovis* ^ko1^H5 parasites by replacement of the *gfp-bsd* selection cassette integrated at the Bb*rad51* locus with the vector, pBb*rad51*wt_comp. pBb*rad51*wt_comp was designed to integrate by double crossover HR (S5 Fig.), replacing the knockout construct with the intact Bb*rad51* wt gene, along with a selection cassette expressing the *hDHFR* gene [33]. CE11 wild-type parasites also were transformed with this vector to control for any effects on parasite viability or sensitivity to MMS caused by manipulation of the Bb*rad51* locus. Importantly, successful complementation by double crossover HR would have resulted in identical Bb*rad51* locus structures in both CE11 and ^ko1^H5 parasites. CE11 parasites incorporated this vector on the first attempt. A similar vector, differing only by the addition of enhanced green fluorescent protein (EGFP) sequences to the 5’ end of the Bb*rad51* open reading frame, also successfully integrated into the CE11 parasite genome without error in two of two attempts, although each crossed over 3’ to the EGFP sequences and failed to incorporate the fluorescent tag. However, HR readily occurred at this locus. In contrast, ^ko1^H5 parasites could not be complemented in this manner in 11 transformation attempts, 7 performed with linearized vector (favoring double crossover homologous recombination) and 4 with circular vector (favoring single-site integration of the entire plasmid). No integration of the transfection plasmid occurred at the Bb*rad51* locus through the intended double-crossover HR, or by single-site integration at the Bb*rad51* locus or ectopically at any other site. Our inability to achieve integration via double-crossover HR in ^ko1^H5 suggests that HR is significantly impaired in the absence of BbRad51, consistent with BbRad51 playing a role in HR-based DNA repair. When tested for MMS sensitivity, CE11/pBb*rad51*wt_comp parasites were indistinguishable from CE11 wild-type (Fig. 4), indicating that modification of the Bb*rad51* locus in this way had not affected the ability of the parasite to repair DNA damage caused by MMS. While we recognize that CE11/pBb*rad51*wt_comp does not represent the complementation of the defect on a ^ko1^H5 background that we had attempted to achieve, we suggest that this represents the probable outcome of successful complementation of the knockout at the Bb*rad51* locus.

### Identification of putative *B. bovis* centromeres

Because we were unable to complement the knockout phenotype through re-integration of the Bb*rad51* gene into the genome, we chose instead to attempt its stable expression from an episomal location. Prior work with an artificial chromosome platform in *P. berghei* [16] suggested this could be a productive approach for expression of exogenous genes in an HR-compromised parasite line. To create one for *B. bovis* we first had to recover native *B. bovis* centromeric and telomeric sequences. Putative *B. bovis* centromeres were identified by focusing on key features common to centromeres in other organisms, including very high percent A+T content, internal repetitive sequences, and a size large enough to mediate attachment of a kinetochore [34, 35]. In this study we searched the genome for A+T-rich regions that are ≥ 2.5 S.D. below the mean G+C content, using a computational window of 1000 bp and length cutoff of ≥ 2 Kbp. Each region so identified was further analyzed by dotplot analysis for internal repetitive structure. Four regions fitting these criteria were identified in the *B. bovis* C9.1 line nuclear genome [36], one per region corresponding to the four *B. bovis* T2Bo chromosomes [25]. The internal repeat structures of the *B. bovis* T2Bo chromosome 2 centromere and of the C9.1 line chromosome 3 centromere were confirmed by dotplot analyses (S6 Fig.). This was done by plotting each sequence against itself, using a 21 nucleotide sliding window (not shown). Each candidate matched well with the predicted centromeric regions of the *B. bovis* T2Bo genome [25], and used by Kawazu and coworkers in constructing the circular centromere-containing plasmid vector, pDHFR-gfp-Bbcent2 [12]. We elected to use the putative centromere from the *B. bovis* C9.1 line chromosome 3 due to its shorter length of 2.3 Kbp, as compared with 3.5- 5 Kbp of the other three, and to ensure full compatibility with derivatives of the *B. bovis* Mexico isolate, such as the C9.1 and CE11 lines.

### Stability of BbACc3

Parasites transformed with BbACc3 (Fig. 5) could be selected with 10 μM pyrimethamine, the approximate IC_90_ for this drug, after approximately three weeks. Parasite growth was somewhat slower than non-transformed parasites, with parasites typically reaching a maximum PPE of approximately 3%. To assess stability of BbACc3, established parasites that had been maintained under pyrimethamine selection for 49 days were split into two. One culture was maintained in the presence of 2 μM pyrimethamine, whereas the second was grown in the absence of drug for 99 days. Each then was grown for 72h in the presence of a range of pyrimethamine concentrations to measure effective IC_50_ values, along with *B. bovis* CE11 and the integrated plasmid line, CE11/pBb*rad51*wt_comp. As seen in Fig. 6, *B. bovis* CE11 had an IC_50_ value for pyrimethamine of approximately 2.5 μM, whereas the line carrying the integrated pBb*rad51*wt_comp plasmid was highly drug-resistant with an IC_50_ of 34 μM. *B. bovis* CE11/BbACc3 that had been maintained under drug selection exhibited an IC_50_ of approximately 24 μM, indicative of high resistance to drug and expression of hDHFR. In contrast, CE11/BbACc3 that had been grown in the absence of drug pressure was indistinguishable from wild type CE11 parasites (IC_50_ = 2.7 μM). These results demonstrate that significant expression of the exogenous gene, hDHFR, occurs from BbACc3, resulting in a ten-fold increase in IC_50_. The drop to wild type levels of drug resistance in parasites grown in the absence of continued selection suggests that either silencing of the hDHFR marker or loss of BbACc3 occurred. We did not pursue identification of the cause, but it is clear that when maintained under drug pressure BbACc3 was maintained and that the hDHFR marker remained actively expressed.

**Fig. 5.**
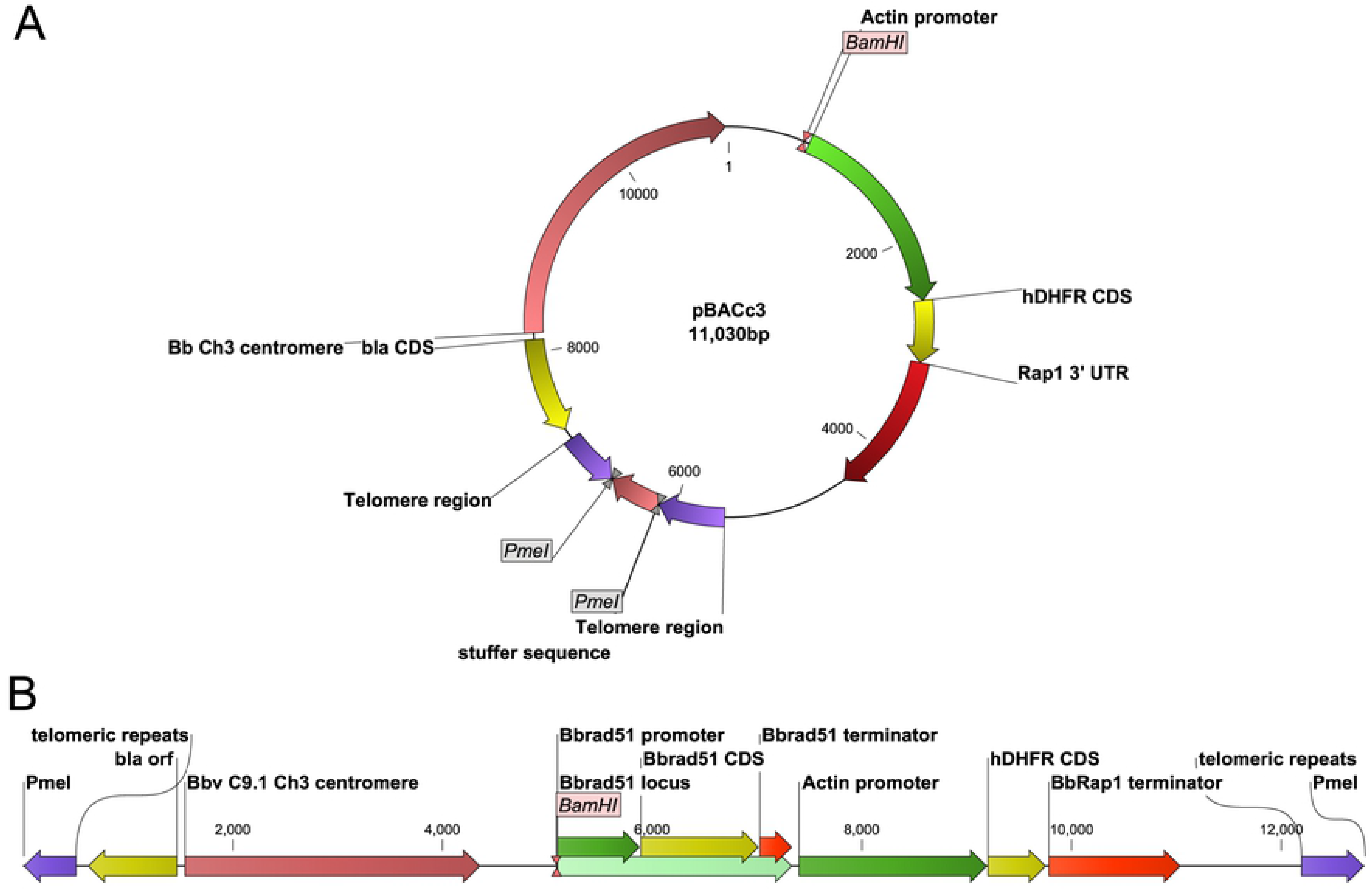
Components and structure of pBbACc3 and pBbACc3_Bb*rad51*wt. **A.** The structure of pBbACc3 in circular plasmid form, prior to removal of stuffer DNA. **B.** Linearized form of BbACc3_Bb*rad51*wt following removal of stuffer DNA by cleavage with PmeI restriction endonuclease. Rap1 3’UTR, *B. bovis rhoptry-associated protein-1* gene 3’ untranslated region; *bla*, β-lactamase coding sequences; CDS, coding sequences; Bb Ch3, *B. bovis* C9.1 line chromosome 3; *hDHFR*, human dihydrofolate reductase coding sequences.

**Fig. 6.**
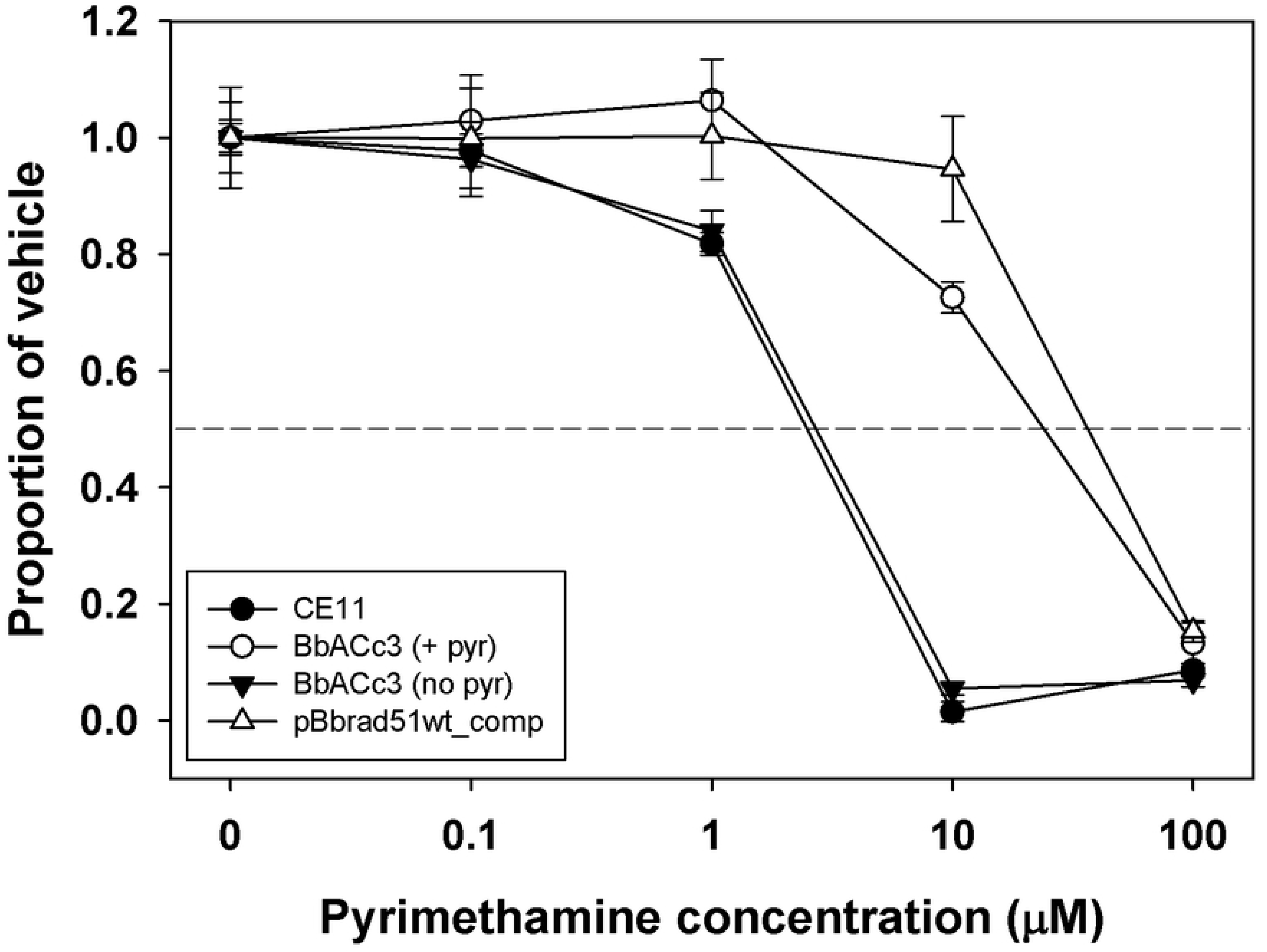
Pyrimethamine-sensitivity in the presence of BbACc3. Sensitivity profiles are shown for *B. bovis* CE11, and CE11 transformed with BbACc3 or pBb*rad51*wt_comp. The presence of either the artificial chromosome, BbACc3, or the integrated construct, pBb*rad51*wt_comp, results in significant resistance to pyrimethamine. The IC_50_ value for each line was approximately 2.5 μM for CE11, 34 μM for CE11/pBb*rad51*wt_comp, 24 μM for CE11/BbACc3 (maintained under constant pyrimethamine pressure; BbACc3 (+pyr)), and 2.7 μM for CE11/BbACc3 three weeks after relief of drug pressure (BbACc3 (no pyr)).

### Adapatation of BbAcC3 derivatives over time

For BbAcC3 and modified derivatives to be used successfully for many purposes would require that this construct behave like a real chromosome, including assembly into chromatin and expansion of telomeric ends for stability. To test for such adaptation over time, we performed partial micrococcal nuclease (MNase) digestion of isolated chromatin. Digestion products were probed by staining with SybrGreen to detect total chromatin, and by Southern blotting to detect BbAcC3_Bb*rad51*-derived sequences. BbAcC3_Bbrad51wt sequences provided a ladder of digestion products with a nucleosomal periodicity of approximately 157-158 bp, matching that of total native chromatin (Fig. 7A) and previously reported values [37], indicative of assembly on nucleosomes. Southern blot probing of isolated plasmid or genomic DNAs, either uncut or cut with informative restriction endonucleases, revealed the expansion of both ends of established BbAcC3 and BbAcC3_Bb*rad51* chromosomes, indicative of telomere lengthening (Fig. 7B).

**Fig. 7.**
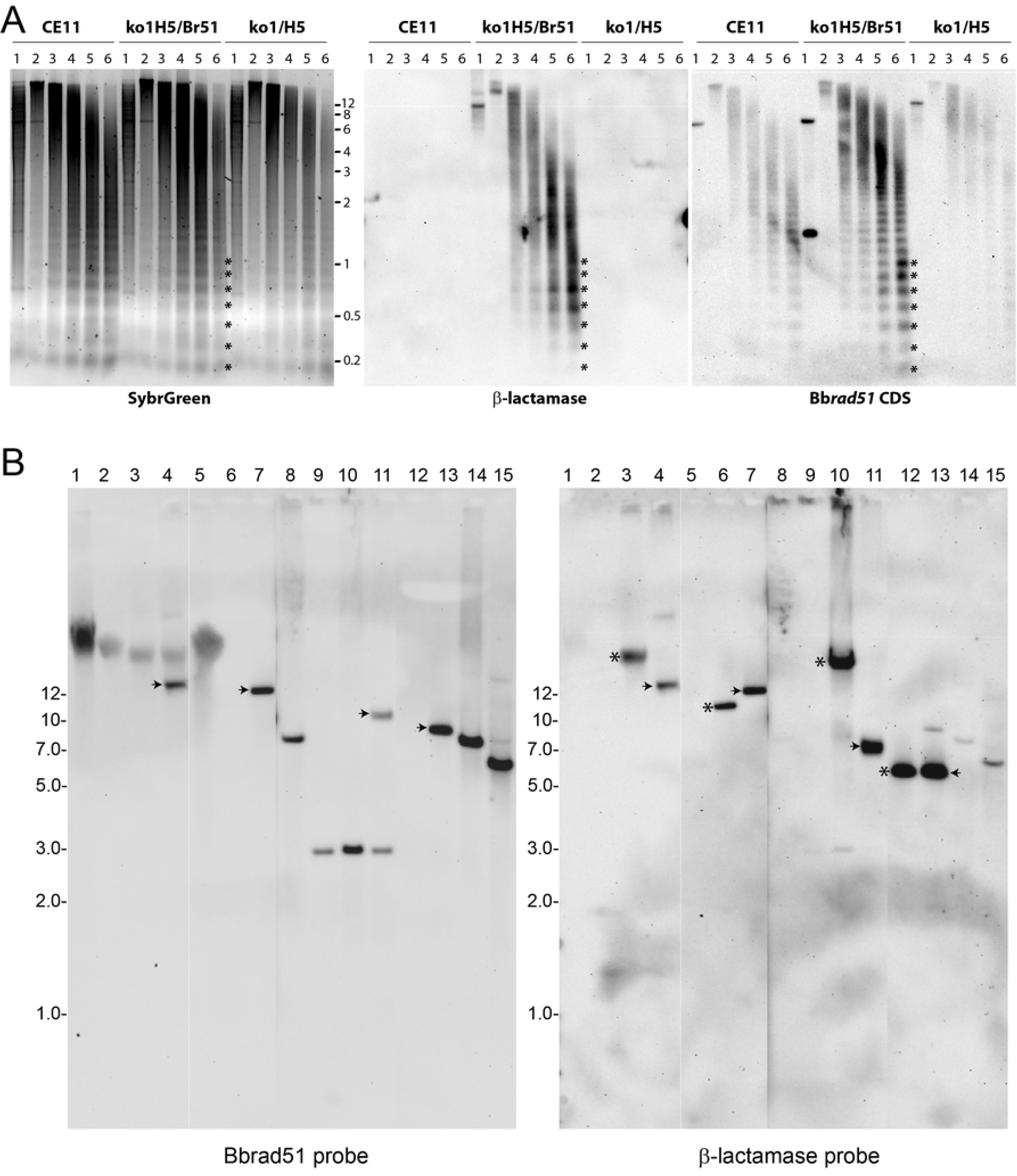
Assembly of BbACc3_Bb*rad51*wt into chromatin. Isolated parasite nuclei were subjected to digestion with titrated concentrations of MNase, then Southern-blotted and probed with different sequences. **A.** (left panel) SybrGreen-stained agarose gel of MNase partially-digested genomic DNAs from *B. bovis* CE11 wild-type (CE11), ^ko1^H5/BbACc3_Bb*rad51*wt (ko1H5/Br51), and ^ko1^H5 Bb*rad51* knockouts (ko1H5). Lanes are: (1) PstI-cut purified DNA, (lane 2) chromatin partially-digested with 0 U, (lane 3) 0.0075 U, (lane 4) 0.015 U, (lane 5) 0.03 U, or (lane 6) 0.06 U of MNase prior to isolation of DNA. Asterisks demonstrate correspondence among the nucleosomal patterns generated from the samples, which form a ladder with 156-159 bp spacing between bands, as previously reported [37]. (center panel) Southern blot of the same gel as in the left panel, probed with β-lactamase sequences to detect the pBluescript backbone of BbACc3_Bb*rad51*wt. (right panel) The same blot stripped and re-probed with Bb*rad51* coding sequences to detect endogenous Bb*rad51* (CE11), both BbACc3_Bb*rad51*wt-provided and non-deleted 3’ sequences (^ko1^H5/Br51), and remaining non-deleted sequences from the 3’ end of B*brad51* (^ko1^H5). **B.** Southern blot probing of plasmids and genomic DNAs from *B. bovis* lines with Bb*rad51* coding sequences (left panel). The same blot was stripped and re-probed with β-lactamase sequences (right panel). Overall telomeric lengthening of BbACc3_Bb*rad51*wt by 2-3 Kbp can be seen by comparing lanes 4 and 7 (both probes, arrowheads). Telomeric lengthening of 1-1.5 Kbp at each end of BbACc3_Bb*rad51*wt can be observed with either probe (compare lanes 11 and 13; arrowheads). Extreme lengthening of the “left” half of BbACc3 (compare lanes 3 and 6, or 10 and 12, asterisks) when using the β-lactamase probe, perhaps due to crossover with a native chromosome. Lanes are: (1) *B. bovis* CE11 wt gDNA; (2) *B. bovis* ^ko1^H5 gDNA; (3) ^ko1^H5/ BbACc3 gDNA (uncut); (4) ^ko1^H5/ BbACc3_Bb*rad51* gDNA (uncut); (5) CE11/ pBb*rad51*wt_comp gDNA (uncut); (6) pBbACc3 plasmid (+ PmeI); (7) pBbACc3_Bb*rad51*wt plasmid (+ PmeI); (8) CE11 gDNA (+ BamHI); (9) ^ko1^H5 gDNA (+ BamHI); (10) ^ko1^H5/ BbACc3 gDNA (+ BamHI); (11) ^ko1^H5/ BbACc3_Bb*rad51*wt gDNA (+ BamHI); (12) pBbACc3 plasmid (+ BamHI, PmeI); (13) pBbACc3_Bb*rad51*wt plasmid (+ BamHI, PmeI); (14) CE11 gDNA (+ EcoRI); (15) CE11/ pBb*rad51*wt_comp gDNA (+ EcoRI). Numbers to the sides of images refer to the positions of double-stranded DNA band size standards, in Kbp.

### Complementation of Bb*rad51* knockout phenotype

Because we were unable to complement the loss of Bb*rad51* by knock-in of the gene via double crossover HR, we attempted complementation with the entire Bb*rad51* locus provided via the BbACc3 platform. CE11 wild type and ^ko1^H5 knockout parasites were transformed by transfection with the artificial chromosome constructs, BbACc3 (as negative control) and BbACc3_Bb*rad51*wt. pBbACc3_Bb*rad51*wt is built upon pBbACc3 but also contains the entire uninterrupted Bb*rad51* locus, including promoter and 3’-termination sequences to facilitate proper regulation (Fig. 5B). ^ko1^H5/ BbAC_Bb*rad51*wt parasites displayed reduced sensitivity to MMS compared with ^ko1^H5 or ^ko1^H5/ BbACc3 parasites, that was not significantly different from wild-type. While not quite reaching wild type levels, this result indicates successful complementation by providing the Bb*rad51* locus on the BbACc3 platform. The contribution of ^ko1^H5/ BbBACc3_Bb*rad51*wt is especially clear when compared with ^ko1^H5/ BbACc3 parasites, which appear to be somewhat less robust overall than ^ko1^H5 (Fig. 8).

**Fig. 8.**
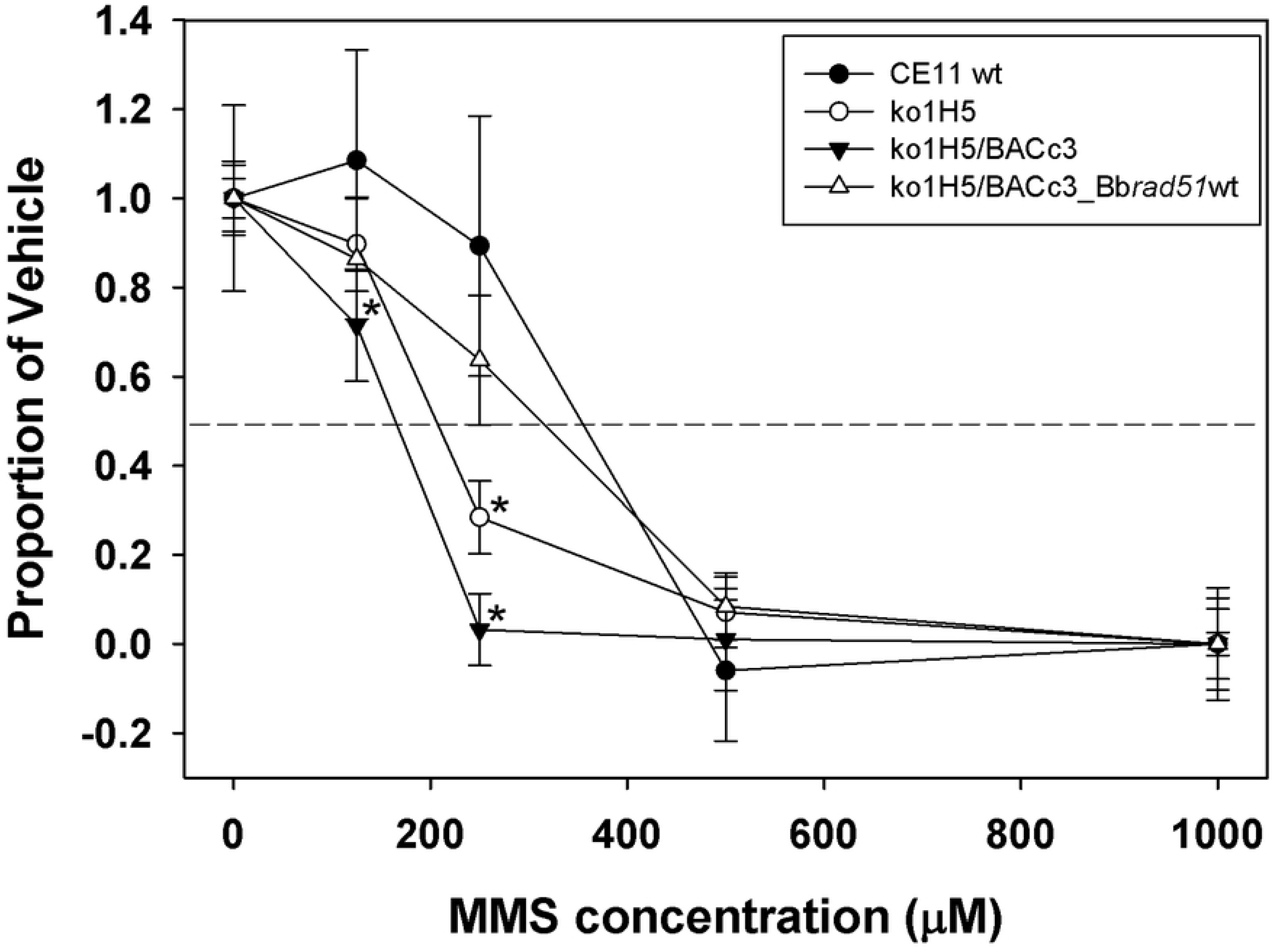
Complementation of the MMS-sensitivity phenotype by Bb*rad51* expressed from the BbACc3 platform. *B. bovis* parasites of the CE11 (wild type), ^ko1^H5 (Bbrad51 knockout), ^ko1^H5/BbACc3_Bb*rad51*wt, and ^ko1^Ht/BbACc3 lines were exposed to varying concentrations of MMS for 90 minutes, followed by drug washout. Parasites then were placed back into culture for 72h, and growth assessed by the SybrGreen method. Growth characteristics were plotted as the growth of each line relative to itself, employing 0 μM as maximal growth and 1000 μM MMS as no growth. Asterisks indicate a difference from the CE11 sample at a significance of p < 0.05. The presence of BbACc3_Bb*rad51*wt complemented ^ko1^H5 sensitivity to 250 μM MMS to a level that was not significantly different from wild-type parasites. The presence of BbACc3 sequences, however, appears to impact somewhat the overall robustness of parasites.

## Discussion

With increasing human population density and environmental encroachment, as well as climate change, many zoonotic diseases are emerging and/or expanding in range, including babesiosis [38–40]. It is thus important that we understand well the biology of these parasites, and prepare to defend against their expansion. Key to understanding the biology of parasites at the molecular level is the ability to manipulate them genetically, something which is not yet well developed for babesial parasites. We have been especially interested in the mechanisms of immune evasion employed by *B. bovis* as a potential “Achilles heel” of this parasite which, if compromised, could lead to its control during infection [41]. At least two mechanisms are used to evade an ongoing immune response: cytoadhesion in the deep vasculature and rapid antigenic variation of the cytoadhesion ligand [6]. Previously, we have shown that antigenic variation relies in large part upon segmental gene conversion (SGC) for modification of the expressed member of the *ves* gene family [42]. With SGC assumed to be a form of homologous recombination, a process thought to be dependent upon Rad51 proteins, we knocked out the Bb*rad51* gene to begin an assessment of its contribution to this phenomenon. Here, we demonstrate that a lack of BbRad51 led to hypersensitivity to MMS, an alkylating agent that methylates adenosine bases [43]. The methyl adducts can result in stalled replication forks and/or single-stranded DNA breaks, both of which can advance to double-stranded DNA breaks, and lead to MMS-hypersensitivity in organisms lacking Rad51 [29–31].

We confirmed in two ways that the MMS-hypersensitive phenotype observed in the Bb*rad51* knockout lines was due to the loss of BbRad51 and not some artifactual secondary effect caused by modification of the genome. First, we integrated pBb*rad51*wt_comp into the Bb*rad51* locus and demonstrated that the phenotype was identical to wt upon exposure to MMS. Secondly, we complemented the phenotype in parasites lacking the Bb*rad51* gene. This was initially attempted with pBb*rad51*wt_comp, which was designed to replace sequences associated with pBb*rad51*ko and reinsert the Bb*rad51* gene back into the genome at its native locus, where it would be under control of its own promoter and terminator elements. However, in 11 attempts, eight performed with linearized vector and three with circular plasmid, no successful transformations were achieved, whereas three out of three attempts were successful in wild-type *B. bovis* CE11 parasites, using linearized vector. We interpret this difference as support for the pivotal importance of Rad51 proteins to HR, including in *B. bovis*. However, while supportive, these results also left the Bb*rad51* gene without complementation to provide formal proof of its involvement. Therefore, we sought to achieve complementation by another means.

In recent years, progress has been made in the ability to genetically modify *B. bovis*, with the development of methods for transient and stable transfection with episomally-maintained plasmids [9, 11, 14, 44–46], and by targeted integration via single- [12, 47] or double-crossover recombination [15, 45]. Although each approach is very useful, each also has its own limitations. Transient transfection provides a short-term phenotype only, does not affect the entire parasite population, and may not accurately reflect gene regulation. Segregation of episomes during mitosis is inaccurate, resulting in a heterogeneous parasite population and plasmid loss from the population over time. Such heterogeneity could compromise the ability to interpret complementation of a phenotype that is not dramatic, such as the MMS-sensitivity phenotype of Bb*rad51* knockout parasites. This problem was partially solved by the inclusion of centromeric sequences into the plasmid which dramatically improved segregation and resulted in sufficient stability for a plasmid to be maintained in the absence of drug selection for two months [12]. However, a circular construct retains the potential to integrate by single-crossover through sequence elements within the plasmid, potentially compromising ongoing studies. We sought to address this issue through the construction of a linear mini-chromosome, BbACc3, containing centromeric and subtelomeric sequences, and telomeric repeats. BbACc3 is manipulated as a plasmid (pBACc3) in *E. coli* for modification and production. It is then linearized for transfection into *B. bovis*, where it is maintained as a linear mini-chromosome with legitimate telomeric ends. Once established, BbACc3_Bb*rad51* was found to assemble with nucleosomes into chromatin (Fig. 7A), and to undergo expansion of the telomeric repeat regions (Fig. 7B). These behaviors suggest that, once established, the BbACc3 platform behaves in a manner consistent with a native chromosome. Here, BbACc3_Bb*rad51*, carrying the Bb*rad51* gene and its native flanking 5’ and 3’ regulatory sequences, was demonstrated to complement the lack of BbRad51 in knockout parasites with regard to MMS-sensitivity. Moreover, complementation occurred at levels not significantly different from wild-type, strongly supporting the notion that BbRad51 contributes to DNA repair and consistent with a possible role in segmental gene conversion. The ability to express sequences in a regulated fashion opens the possibility of using BbACc3 as a platform to complement knockout of essential genes, or to perform studies of promoter structure and functional interactions. Moreover, constructs carrying exogenous genes could be used to engender transmission-blocking immunity by including vector proteins, to immunize against other co-transmitted parasites by inclusion of appropriate vaccine genes, or other similar applications.

In the course of these studies we found it useful to construct and employ an artificial chromosome, BbACc3, as a platform for the reintroduction of the Bb*rad51* gene into *B. bovis*. This allowed us to circumvent the need for homologous recombination to reintegrate Bb*rad51* sequences into the genome in order to acquire a uniform population. Significantly, BbACc3_Bb*rad51*wt enabled the complementation of the Bb*rad51* knockout phenotype at near-normal levels in response to MMS challenge. Once established, the BbACc3 platform behaved like a native chromosome, forming chromatin and lengthening its telomeres. Despite these positive attributes, a cautionary note must also be provided. Southern blot results suggest that within CE11 wild-type parasites, the major parasite population to grow up during selection appears to have undergone a crossover event between BbACc3 and one of the native chromosomes (Fig. 7B). This appears to have occurred through subtelomeric or telomeric repeat sequences at the “left” end near the β-lactamase coding sequences, resulting in significant expansion of that end of BbACc3, and an overall much longer artificial chromosome than in ^ko1^H5/BbACc3_Bb*rad51*wt, even after expansion of its telomeric repeats. An event of this nature would always be a potential hazard in any HR-competent parasite line. A useful improvement of BbACc3 could be made through the inclusion of replication origin sequences that might improve the ease of initial establishment of the chromosome, which in these experiments was no faster than selection of double crossover integration mutants. Thus, while artificial chromosome technology holds considerable promise for certain applications and was invaluable in this study, it will require optimization to meet its full potential. Regardless, BbACc3 provides a new option in the toolkit for genetic manipulation of *B. bovis* and perhaps other babesial parasites where its centromeric and regulatory sequences may function, and will be made freely available to the scientific community.

From these studies we conclude that Bb*rad51* encodes the *B. bovis* Rad51 ortholog, and that this protein is non-essential to the parasite for in vitro survival and growth. It remains to be determined whether the parasite could similarly survive during infection of a host without BbRad51. Further, we provide evidence that the encoded protein is involved in recovery from DNA damage, consistent with known functions of Rad51 proteins in other systems [48]. However, any connection of BbRad51 with SGC specifically remains to be established.

## Methods

### Parasites and culture conditions

*B. bovis* parasites of the CE11 clonal line were used in experiments [7]. Parasites were grown under microaerophilous stationary phase culture conditions, with an atmosphere of 90:5:5 nitrogen: oxygen: carbon dioxide [49, 50]. Defibrinated bovine blood products were obtained locally from Holstein cows (approved by University of Florida Institutional Animal Care and Use Committee, protocol #201102216) or commercially (Hemostat Laboratories; Dixon, CA). Parasite sensitivity to pyrimethamine was determined as described [51]. Transformed parasites were maintained in medium containing 2.0 μM pyrimethamine.

### Validation of Bb*rad51* gene and BbRad51 identity

Identification of BbRad51 (accession no. XP_001609877) was achieved by reciprocal Psi-BLAST searches of the *B. bovis* T2Bo genome [25], using *S. cerevisiae* Rad51 (accession no. CAA45563) to initiate the query. Alignments used sequences publicly available through Genbank, EuPathDB database (http://eupathdb.org/eupathdb/), and/or Wellcome- Sanger Institute pathogens ftp site (ftp://ftp.sanger.ac.uk/pub/pathogens/). Annotation of Bb*rad51* gene structure was confirmed at both the genome (GeneID: 5478106) and transcript (XM_001609827.1) levels as described below. Virtual translation and routine manipulations performed with CLC-Bio Main Workbench (CLC-Bio, Arrhenius, Denmark). Conserved Domains search was performed through the National Center for Biotechnology CDD webserver (http://www.ncbi.nlm.nih.gov/Structure/cdd/cdd.shtml) [52, 53]. Three-dimensional structural modeling was performed through the Robetta webserver (http://robetta.bakerlab.org/) [22, 54]. Structural superimpositions and graphics were created with the Chimera software package of the Resource for Biocomputing, Visualization, and Informatics at the University of California, San Francisco (www.cgl.ucsf.edu/chimera/), using the Matchmaker algorithm [24].

### Validation of Bb*rad51* gene structure

*B. bovis* gDNAs were isolated as described [55, 56], but with prior ammonium chloride lysis of the erythrocytes [57]. Alternatively, QIAmp Mini Spin Columns (Qiagen) were sometimes used, following manufacturer’s instructions. For RNA extractions, cultures were grown in erythrocytes from which host leukocytes were removed by filtration through Whatman CF-11 cellulose (GE Healthcare Life Sciences; Pittsburgh, PA) [58]. When needed, cultures were grown to elevated levels of percent parasitized erythrocytes by “dilution enrichment” [59]. Packed erythrocytes were emulsified with either TRIzol reagent (Invitrogen; Waltham, MA) or RiboZol reagent (Amresco; Solon, OH), extracted twice with chloroform and precipitated from 2-propanol. A 2244 bp segment containing the 1134 bp Bb*rad51* gene and its 5’ and 3’ intergenic regions was amplified from *B. bovis* C9.1 gDNA by polymerase chain reaction (PCR) with Phusion High-Fidelity DNA Polymerase (New England BioLabs; Beverley, MA) using primers EAM6 and EAM18. This segment was cloned into pCR2.1-TOPO-TA (Invitrogen) to create plasmid pBbRad51. Bb*rad51* structure was confirmed by PCR amplification of the 1134 bp gene within the 2244 bp amplicon, using flanking primers EAM6 and EAM18. The amplicon was cloned into pCR2.1-TOPO-TA (Invitrogen) and sequenced through the University of Florida Sanger Sequencing Core by primer-walking. To characterize Bb*rad51* transcripts, cDNA was made from *B. bovis* C9.1 line total RNA, using M-MuLV Reverse Transcriptase (New England Biolabs; Beverley, MA) and oligo-d(T) primer. Full-length coding sequences were obtained by PCR amplification of cDNA with primers EAM50 and EAM51. The amplicon was cloned into pCR2.1-TOPO-TA and sequenced. To obtain non-coding transcript sequences, RNA Ligase-mediated rapid amplification of cDNA ends (RLM-RACE) was performed, using the FirstChoice RLM-RACE kit (Ambion; Waltham, MA). Tobacco acid pyrophosphatase was used to remove the 5’-cap structure for adapter addition, and reverse transcription using M-MuLV and random decamers. Nested PCR amplification of 5’-untranslated sequences was performed using primers EAM2 and 5’-RACE outer, and EAM4 and 5’-RACE inner primers. Products were cloned into pCR-TOPO-Blunt and sequenced. The 3’-untranslated sequences were obtained by addition of the 3’-RACE adapter, and nested PCR amplification with primers EAM1 and 3’-RACE outer, and EAM3 and 3’- RACE inner. Products were directly cloned into pCR-TOPO-Blunt and sequenced.

### Bb*rad51* knockout plasmid assembly

The Bb*rad51* gene was knocked out by a double crossover homologous recombination strategy [60], with plasmid pRad51KO (Fig. 2A). Plasmid pRad51KO was designed to replace all but the 3’-terminal 388 bp of Bb*rad51* coding sequences with a selectable *gfp-bsd* marker (a gift from C.E. Suarez). This region was retained because of the close proximity (320 bp) of Bb*rad51* to the gene downstream encoding a conserved mechanosensitive channel-like protein, XP_001609876. To assemble pRad51KO, the *EF1α* gene “B” hemi-promoter for driving the *bsd* selectable marker was PCR-amplified from *B. bovis* CE11 line gDNA with primers EAM42 and EAM43. Bb*rad51* 3’-end targeting sequences were amplified from plasmid pBbRAD51, using primers EAM49 and EAM56. The *EF1α*-B segment and Bb*rad51* 3’ targeting sequences were combined by crossover PCR [61] with EAM43 and EAM56 to form fragment 3’-*rad51-EF1α*-B. The 92 bp *B. bovis* β-tubulin terminator region (T3) was amplified from plasmid pDS-*bsd* [11], using primers EAM45 and EAM46. The *gfp-bsd* fusion gene from plasmid pGFP-Bsd [14], was combined with the T3 fragment by crossover PCR, using primers EAM44 and EAM46 and including the T3 fragment in the reaction. The Bb*rad51* upstream targeting sequence was added to the *gfp-bsd*-T3 fragment by crossover PCR with primers EAM44 and EAM47, and including pBbRad51 in the reaction, to form fragment *gfp-bsd*-T3-5’-*rad51*. The two fragments were combined by crossover PCR, using primers EAM56 and EAM47 and inserted into pBluescript II-KS(+). A single missense mutation was identified within the *bsd* sequence. This was corrected by PCR-amplification of the two halves of the insert with EAM61 and EAM47, and EAM62 and EAM56. The two fragments were fused by crossover PCR, using only EAM47 and EAM56, and reinserted into pBluescript II-KS(+). The corrected plasmid, pRAD51KO, was transformed into *Escherichia coli* DH5α. Proper construction was confirmed by full sequencing, and the corrected plasmid used for subsequent plasmid production.

### Validation of Bb*rad51* knock-out

Parasite populations recovered initially were screened directly with the Phusion Blood Direct PCR Kit (Thermo Scientific), using primers EAM75 and EAM77 within the coding sequences of the immediately flanking genes. Non-mutated parasites generated a band of 2.4 Kbp, whereas a proper knockout yielded a band of 3.4 Kbp. RT-PCR with primers EAM50 and EAM51 confirmed the lack of Bb*rad51* transcripts. Southern blots were performed as described [42]. DNA fragments, alkaline-transferred to nylon membranes (Amersham Hybond-N^+^, GE Healthcare), were cross-linked with 50 mJ ultraviolet in a GS Gene Linker UV chamber (BioRad; Hercules, CA). Oligonucleotide probes were end-labeled with γ-[^32^P]ATP, using polynucleotide kinase.

### Parasite growth assays

Parasite growth was assayed by counting Giemsa-stained smears, with samples collected at 0, 24, and 48h growth (approximately 0, 3 and 6 cell cycles [62]). Alternatively, in some experiments a DNA-based SYBR Green I method was performed, essentially as described [63, 64], on parasites grown in bovine erythrocytes depleted of leukocytes [58]. For measuring sensitivity to pyrimethamine, parasites were grown in complete medium containing serial 10-fold dilutions of pyrimethamine for 72 hours, at concentrations from 0 to 100 μM. For experiments involving MMS, parasites were exposed at 2% ppe to various concentrations of MMS (diluted in complete medium) for 90 minutes at room temperature, then were washed two times at 6000 x g.min at room temperature with 1x VYM buffer [65] to remove MMS. Parasites then were resuspended in complete medium, and placed under normal culture conditions. In initial experiments, cells were diluted 1/10 into a 10% packed cell volume suspension of untreated uninfected erythrocytes, and were placed back into culture. Smears were made at 0, 24, 48, and 72 hours post-treatment, and Giemsa-stained for microscopic counting of percent parasitized erythrocytes. After validation of effect, subsequent experiments were performed using the SybrGreen method, with starting values of 0.2% parasitized erythrocytes, and terminal collection at 72 hours only. An initial titration experiment with *B. bovis* CE11 established 125 μM to 1000 μM as the informative range.

### Assembly of complementation constructs

Two different complementation strategies were used in this project. (i) In the first strategy, plasmid pBb*rad51*wt_comp (S5 Fig.) was engineered to replace all sequences associated with the knockout plasmid by double crossover homologous recombination. The goal was to replace them with the wild-type Bb*rad51* locus, concomitant with integration of a human dihydrofolate reductase (hDHFR) expression cassette, to enable selection of transformants with pyrimethamine [51]. pBb*rad51*wt_comp was made by PCR-amplifying Bb*rad51* 3’ terminator sequences from *B. bovis* C9.1 line gDNA with primers DA255 + DA256. The hDHFR selection cassette was moved from pBACc3 by amplification with primers DA254 + DA253. pBluescript-SK(+) was linearized with XhoI + HindIII restriction endonucleases, and the three sequences were assembled simultaneously, using InFusion reagents (Clontech). The Bb*rad51* locus, including complete promoter through immediate 3’ sequences, was amplified from *B. bovis* C9.1 line gDNA with primers DA248 + DA252. The Bb*rad51* locus was inserted, using InFusion reagents, into the intermediate construct that had been opened with XbaI + HindIII. (ii) The second strategy was to reintroduce the Bb*rad51* locus in the context of the BbACc3 artificial chromosome, which did not require reintegration into the genome but allowed expression controlled by autologous regulatory elements. Construction of pBACc3 (circular plasmid form) was begun with destruction of the NotI site (nt 669-676) within the multiple cloning site (mcs) of pBluescript-KS(+), by a fill-in reaction with T4 DNA polymerase and blunt ligation [66]. This product was partially digested with RcaI, and amplified by inverse-PCR with Phusion Hotstart II polymerase (New England Biolabs; Beverley, MA), using primers DA162 and DA163 [67]. Upon circularization of the amplicon nt 2888-2940 were deleted, replacing the RcaI site at position 2881-2886 with a new NotI site, and creating the 2920 bp intermediate plasmid, pBS2. *B. bovis* C9.1 line telomeric sequences were inserted into the remaining RcaI site at nt 1873-1878 (original numbering), as follows: *B. bovis* C9.1 line genomic DNA (gDNA) was PCR-amplified with Phusion polymerase (New England Biolabs; Beverley, MA), using primers specific to unique sequences at nt 2,592,052-2,592,072 (DA170) and 948-923 (DA171) of the *B. bovis* T2Bo chromosome 3, coupled with primers DA172 or DA173, respectively. The identification of centromeric sequences is provided in S6 Fig.. The 5’ halves of DA170 and DA171 were complementary to vector sequences flanking the remaining RcaI site. The 3’ halves of primers DA172 and DA173 each represented three telomeric repeats. The 3’-most repeat of each contained an extra T, an infrequent imperfection in the telomeric repeats of *B. bovis* (T2Bo isolate reference genome, accession no. AAXT01000001.1) [25], whereas the 5’ halves overlapped with *Anaplasma marginale* genomic sequences. A 483 bp stuffer fragment, corresponding to nucleotides 145014-145482 of the *A. marginale* Florida isolate genome (accession no. NC_012026), then was amplified with primers DA152 and DA153, and a two-step crossover PCR approach was used to assemble all three amplicons into one fragment containing PmeI restriction sites flanking the stuffer [61]. The fused amplicon was inserted into the RcaI site, using InFusion reagents (Clontech Laboratories, Inc.; Mountain View, CA). Putative centromeres were identified in the *B. bovis* C9.1 line genome (ftp://ftp.sanger.ac.uk/pub/pathogens/Babesia/). Artemis v.16.0.0 (http://www.sanger.ac.uk/science/tools/artemis) was used to observe sequence %GC content, identifying stretches > 2.5 standard deviations below the mean in a running window of 1000 nt [68]. Sequences were considered candidates when the AT-rich segment was 2 Kbp or longer and possessed no annotated open reading frames. Candidate sequences were further analyzed by dot-plot to identify those containing significant internal repetitive sequence structure (S6 Fig.). A single candidate was identified from chromosome 3, corresponding to position 2922284-2925108. Centromeric sequences were amplified with primers DA174 and DA175, and inserted into the NotI site by ligation with T4 DNA ligase. The hDHFR selection cassette was moved from the plasmid pDHFR-gfp-Bbcent2 (a gift from Shin-Ichiro Kawazu [12]) into the intermediate construct, in multiple steps. PCR-amplification of the hDHFR coding sequences with DA164 + DA165, the *B. bovis* actin promoter with DA166 + DA167, and the Rap1 gene 3’ terminator sequences with DA168 + DA169 was used to recover these segments. A two-step crossover PCR was used to combine hDHFR coding sequence with promoter and terminator [61], simultaneously removing all HindIII, EcoRV, and PstI sites. The full-length cassette first was inserted into the SalI site of pBS2. The complete cassette then was removed with KpnI and PstI, and integrated into the artificial chromosome construct, using Gibson Assembly reagent mix (New England Biolabs; Beverley, MA) [69], yielding pBACc3. The final pBACc3 basic vector construct is 11,030 bp (Fig. 5A). To make BbACc3_Bb*rad51*wt, the entire Bb*rad51* locus was amplified from pBb*rad51*wt_comp with primers DA248 + DA281, and inserted with NEBuilder reagents (New England Biolabs) into pBACc3 opened with BamHI and PstI. All added sequences and modified regions of the plasmid were fully Sanger-sequenced to confirm proper construction. Constructs were transfected into *E. coli*, using strains DH5α (Invitrogen), NEB10β (New England Biolabs), or Stellar (Clontech), by electroporation. For transfection, pBACc3_Bb*rad51*wt was linearized with PmeI to release the stuffer sequence and expose the telomeric ends (Fig. 5B). pBACc3 will be made available upon request.

### Parasite transformation, and maintenance of BbACc3

Parasites were washed into cytomix buffer [70], then transfected with knockout or complementation vectors dissolved in cytomix. Transfection was achieved by electroporation with 10 pmol of DNA, under conditions of 1.25 kV, 25 μF, and 200 Ω resistance, as described [11]. Prior to transfection, pRad51ko and pBbrad51wt_comp were linearized with NotI. pBACc3_Bb*rad51*wt linearized with PmeI, releasing the *A. marginale* “stuffer” DNA and generating linear BbACc3_Bb*rad51*wt with telomeric ends. It was found not to be necessary to purify BbACc3 away from cleaved stuffer DNA prior to transfection. In some experiments parasites were transfected with pBACc3 as supercoiled plasmid. Following transfection, parasites were placed into culture for 24 h in the absence of drug selection. To effect selection, pyrimethamine then was added and maintained at a concentration of 2 μM (the approximate IC_90_ concentration), beginning 24h post-transfection. In some experiments, drug pressure was removed for 21 days (approximately ≥ 60 cell cycles) from already-transformed parasites, then was reapplied at varying concentrations to assess for loss of drug-resistance.

### Observation of chromosome maintenance and chromatin assembly

Maintenance of BbACc3_Bb*rad51*wt as a linear chromosomal element was confirmed by isolation of genomic DNA [55], with resolution of uncut and restriction-digested DNAs on a 0.7% agarose gel. DNAs were alkaline-transferred to nylon membranes as described [42], and hybridized with probes to Bb*rad51*, β-lactamase, centromeric, and hDHFR sequences. Probes were generated by PCR amplification of pBACc3_Bb*rad51*wt sequences with primers EAM8 + EAM11, DA290 + DA291, DA189 + DA190, or DA164 + DA165, etc., respectively. One μg of each amplicon was labeled with digoxigenin, and detected with anti-digoxigenin-HRP antibodies and CSPD substrate, according to supplier’s instructions (Sigma Chemical; St. Louis, MO). Blots were imaged with a FluorChemR instrument (Protein Simple; San Jose, CA) by detection of luminescence. The stability of telomeric end lengths was tested by assessing the lengths of BamHI restriction fragments of PmeI-linearized BbACc3 constructs used to transfect, and BbACc3 constructs recovered after establishment of transformed parasites in culture. gDNAs were cleaved with BamHI, and probed with the above-mentioned probes. pBACc3 digested with PmeI to release stuffer sequences was used as control. Probes were labeled with 11-dUTP-digoxigenin, following manufacturer’s instructions, and detected as described above. To observe assembly of BbACc3 and BbACc3_Bbrad51wt into chromatin, nuclei were isolated from transformed parasites, the chromatin partially digested with micrococcal nuclease (MNase), and the fragments isolated as described [37]. To observe overall nucleosomal assembly and spacing, digestion products were stained in-gel with SybrGold. Similar gels were also alkaline-blotted to nylon membranes and probed by Southern blotting, as described above. Alkali-labile probes were used to facilitate stripping of blots and re-probing.

## Acknowledgments

The authors thank Carlos Suarez and Shen-ichi Kawazu for providing plasmids pGFP-Bsd and pDHFR-gfp-Bbcent2, respectively, and Kevin Brown and Linda Bloom for providing critical early comments. We are indebted to Allison Vansickle for assistance with bovine blood collection and maintenance of in vitro cultures.

## Supporting Information Captions

**S1 Fig.. Similarity of *B. bovis* RecA/RadA/Rad51 superfamily-related proteins to known or putatively annotated Rad51 proteins from other species.** Those proteins for which there is experimental support for catalysis of canonical Rad51 functions are indicated in blue in Fig. 1. The Walker A and Walker B motifs are indicated here by red and blue box overlays, respectively. This alignment provided the basis for the phylogenetic tree shown in Fig. 1A.

**S2 Fig.. RT-PCR amplification of Bb*rad51* and *gfp-bsd* transcripts.** Total RNAs were isolated from *B. bovis* CE11 subclones B8, C2, and C5, the initial uncloned CE11ΔBb*rad51* knockout and three clonal lines derived from it (^ko1^C3, ^ko1^H5, and ^ko1^H6), and lines CE11Δrad51ko2 and CE11Δrad51ko3. **A.** cDNAs made with oligo[dT] primer were amplified with primers EAM8 and EAM11 [11, 51] for detection of Bb*rad51* transcripts. **B.** cDNAs were amplified with XW119 and XW121 [11, 51] for detection of *gfp-bsd* transcripts. Results demonstrate Bb*rad51* transcription by wild type lines but not by knockouts. Conversely, *gfp-bsd* transcripts are present in knockout but not wild type lines.

**S3 Fig.. Southern blot analysis of Bb*rad51* locus. A.** Schematic diagram of the Bb*rad51* locus in *B. bovis* CE11 wild type (bottom) and knockout (top) parasites, and the locations of restriction endonuclease and primer binding sites. The tables provide the anticipated sizes (in bp) of specific fragments detected with the indicated probes, based upon the genome sequence. **B.** Southern blots of *B. bovis* CE11 wild type, and the initial CE11ΔBb*rad51* knockout parasite population both early and late (12 months later) in selection prior to cloning (left panel), after probing with Bb*rad51*-specific oligonucleotide probe, EAM10. Initial *B. bovis* CE11ΔBb*rad51* clonal lines ^ko1^C3, ^ko1^H5, and ^ko1^H6 are shown in the right panel. **C.** The same blots shown in panel B are shown after being stripped and re-probed with oligonucleotide DA101R, specific for *gfp* sequences. The numbers above the bands indicate the sizes of the bands in bp.

**S4 Fig.. Titration of *B. bovis* CE11 wild type parasite sensitivity to MMS.** In initial experiments to determine useful concentrations of MMS to observe phenotype, parasites were exposed to varying concentrations of MMS for 90 minutes at room temperature, then washed and placed back into culture for 72h. Smears were made at 0, 48, and 72h post-treatment, fixed and stained with Giemsa stain, and manually counted by microscopy in a blinded fashion. Growth is indicated as percent parasitized erythrocytes (PPE). This experiment indicated that a range from 250 μM to 1000 μM MMS provided useful results.

**S5 Fig.. Structure of the pB*brad51*wt_comp double crossover complementation plasmid.** This plasmid was designed to integrate into the genome by double-crossover homologous recombination via sequences upstream and downstream to the Bb*rad51* coding sequences. In doing so, the strategy was to recreate the Bb*rad51* locus, but with a short 3’-untranslated region for regulation, followed by the hDHFR selection cassette. The plasmid was introduced into parasites after NotI linearization, and selected for growth in the presence of pyrimethamine. This occurred as intended in three of three attempts with wild type parasites, but failed in 11 attempts with *B. bovis* ^ko1^H5 parasites.

**S6 Fig.. Identification of the *B. bovis* C9.1 chromosome 2 centromere. A.** *B. bovis* T2Bo isolate genomic sequences [25] were scanned with the “GC Content (%) with 2.5 SD Cutoff” subroutine of Artemis v. 16.0.0 [68], using a sliding window of 1000 bp. Stretches of sequence > 2.5 standard deviations below the mean G+C content, and ≥ 2 Kbp in length, were identified. Shown is the region of chromosome 3 identified by this strategy. Sequence comprising nucleotides 2922285-2925108 of the *B. bovis* C9.1 line genome [36], corresponding to nucleotides 29912-32736 of the *B. bovis* T2Bo isolate chromosome 3 [25], were recovered by PCR and used in the construction of pBbACc3. **B.** Dot-plots of the putative *B. bovis* C9.1 line chromosome 3 (left plot) and T2Bo isolate chromosome 2 (right plot) centromeres against themselves. The internal repeat structure of each is apparent from the plots, with the chromosome 2 centromere having a larger major repeat domain. The darkness of spots indicates the degree of similarity, with the dark blue diagonal lines indicating identity. Values within the range from 40-100% identity is shown. When plotted against one another there is no evidence for any specific sequence relationship (not shown).

